# *Staphylococcus aureus* does not synthesize arginine from proline under physiological conditions

**DOI:** 10.1101/2022.01.12.476138

**Authors:** Bohyun Jeong, Majid Ali Shah, Eunjung Roh, Kyeongkyu Kim, Indal Park, Taeok Bae

**Affiliations:** Department of Microbiology, Research Institute for Microbial Resistance, Kosin University College of Medicine, Busan, Korea; Department of Microbiology and Immunology, Indiana University School of Medicine-Northwest, Gary, Indiana, 46408, USA; Korea Crop Protection Division, National Institute of Agricultural Sciences, Rural Development Administration, Wanju-gun, Korea; Department of Molecular Cell Biology, Sungkyunkwan University School of Medicine, SMC, Suwon, Korea

## Abstract

The Gram-positive pathogen *Staphylococcus aureus* is the only bacterium known to synthesize arginine from proline via the arginine-proline interconversion pathway, despite having genes for the well-conserved glutamate pathway. Since the proline-arginine interconversion pathway is repressed by CcpA-mediated carbon catabolite repression (CCR), CCR has been attributed to the arginine auxotrophy of *S. aureus*. Using ribose as a secondary carbon source, here, we demonstrate that *S. aureus* arginine auxotrophy is not due to CCR but due to the inadequate concentration of proline degradation product. Proline is degraded by proline dehydrogenase (PutA) into pyrroline-5-carboxylate (P5C). Although the PutA expression was fully induced by ribose, the P5C concentration remained insufficient to support arginine synthesis because P5C was constantly consumed by the P5C reductase ProC. When the P5C concentration was artificially increased by either PutA overexpression or *proC*-deletion, *S. aureus* could synthesize arginine from proline regardless of carbon source. In contrast, when the P5C concentration was reduced by overexpression of *proC*, it inhibited the growth of the *ccpA*-deletion mutant without arginine. Intriguingly, the ectopic expression of the glutamate pathway enzymes converted *S. aureus* into arginine prototroph. In an animal experiment, the arginine-proline interconversion pathway was not required for the survival of *S. aureus*. Based on these results, we concluded that *S. aureus* does not synthesize arginine from proline under physiological conditions. We also propose that arginine auxotrophy of *S. aureus* is not due to the CcpA-mediated CCR but due to the inactivity of the conserved glutamate pathway.

**Importance:** *Staphylococcus aureus* is a versatile Gram-positive human pathogen infecting various human organs. The bacterium’s versatility is partly due to efficient metabolic regulation via the carbon catabolite repression system (CCR). *S. aureus* is known to interconvert proline and arginine, and CCR represses the synthesis of both amino acids. However, when CCR is released by a non-preferred carbon source, *S. aureus* can synthesize proline but not arginine. In this study, we show that, in *S. aureus*, the intracellular concentration of pyrroline-5-carboxylate (P5C), the degradation product of proline and the substrate of proline synthesis, is too low to synthesize arginine from proline. These results call into question the notion that *S. aureus* synthesizes arginine from proline.

## Introduction

*Staphylococcus aureus* is a Gram-positive pathogenic bacterium causing a wide range of infections in humans (1). *S. aureus* genomes contain genes for the synthesis of all 20 amino acids (2). Intriguingly, however, the bacterium is phenotypically auxotrophic to several amino acids, including arginine and proline. The arginine and proline auxotrophy of *S. aureus* is attributed to the CcpA-mediated carbon catabolite repression (CCR) (3, 4).

CcpA is a transcription factor of the LacI/GalI family and plays a diverse role from CCR, nitrogen metabolism, and virulence (5, 6). In the CCR, when a preferred carbon source such as glucose is present, CcpA represses the transcription of the genes necessary for utilizing non-preferred carbon sources, including ribose, arabinose, and sorbitol, allowing the bacterium to avoid unnecessary transcription and to optimize energy usage (7). The CcpA-binding site is known as catabolite response element (CRE) (8), and it consists of the following consensus sequence: WWTGNAARCGNWWNCAWW (W = A or T; R = A or G, N = any). Typically, the CRE-binding of CcpA requires the corepressor protein HPr (histidine protein) (9).

In *S. aureus*, the synthesis pathways for arginine and proline are interconnected: arginine is synthesized from proline, while proline is synthesized from arginine (Fig. 1A) (3, 4) (Fig. 1). In fact, *S. aureus* is the only bacterium known to synthesize arginine from proline (4). In the arginine-proline interconversion pathway, CcpA represses almost all steps except for the final proline synthesis by ProC (3, 4)(Fig. 1A). Intriguingly, in addition to the arginine-proline interconversion pathway, *S. aureus* also contains the well-conserved glutamate pathway for arginine synthesis. This pathway converges to the arginine-proline interconversion pathway at ornithine (Fig. 1A). Intriguingly, the *argDCJB* operon, encoding the unique steps of the glutamate pathway, is not transcribed in *S. aureus* even under 44 different growth conditions (4, 10).

**Fig. 1.**
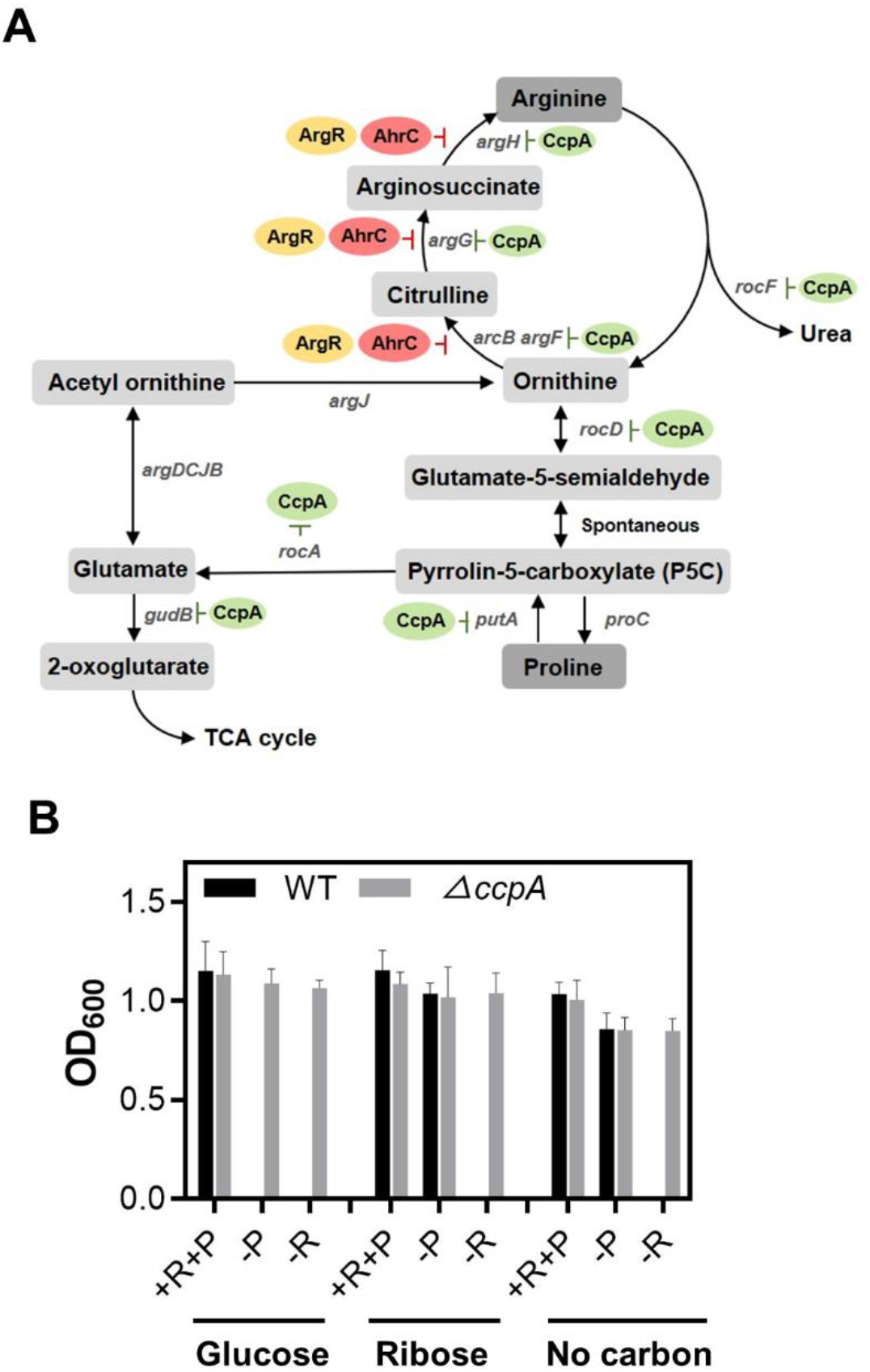
*S. aureus* remains an arginine auxotroph regardless of carbon sources. (A) Metabolic pathways for arginine and proline in *S. aureus*. The steps repressed by CcpA (green) and arginine repressors (yellow and red) are indicated. (B) Carbon source effect on arginine and proline auxotrophy of *S. aureus*. *S. aureus* wild-type (WT) and the *ccpA* mutant (Δ*ccpA*) were grown at 37C° overnight in CDM containing glucose or ribose or no carbon source. Bacterial growth was measured by optical density at 600 nm (OD_600_). -, absence; +, presence; R, arginine; P, proline.

At the center of the intertwined metabolic pathways is pyrroline-5-carboxylate (P5C). P5C is a degradation product of ornithine and proline and can be used for the synthesis of three amino acids: arginine, proline, and glutamate (Fig. 1A). In particular, the P5C-glutamate conversion allows *S. aureus* to use arginine and proline as alternative carbon sources in the absence of glucose because glutamate can be further converted into 2-oxoglutarate, an intermediate of the TCA cycle, by GudB (Fig. 1A) (11). Not surprisingly, these P5C to 2-oxoglutarate conversion steps (i.e., the productions of P5C dehydrogenase (RocA) and GudB) are also repressed by CcpA (Fig. 1A).

Since the arginine-proline interconversion pathway is repressed by CcpA, *S. aureus* is expected to become a prototroph for both arginine and proline when *ccpA* is knocked out. Indeed, the *ccpA*-knock out mutant can grow in the absence of arginine or proline (3, 4). Surprisingly, however, when the CcpA-mediated CCR is abolished by using secondary carbon sources such as ribose as a sole carbon source, although *S. aureus* became a prototroph for proline (3), it still remained an arginine auxotroph(4). Also, by an experiment with ^13^C-labeled amino acids, Halsey et al. clearly demonstrated that proline is not converted into arginine even in the absence of glucose (11). Therefore, it is questionable whether *S. aureus* synthesizes arginine from proline under physiological conditions (i.e., without gene knockout).

In this study, using ribose as a secondary carbon source, we wanted to understand the molecular mechanism behind the arginine auxotrophy in the absence of CCR and investigate whether the *argDCJB* operon is functional when ectopically expressed. We found that the insufficient concentration of proline degradation product P5C, not CCR, is responsible for the arginine auxotrophy of *S. aureus*, and the *argDCJB* operon is functional once transcribed.

## Results

### The CCR of the arginine synthesis is not affected by the carbon source

Previously, in the chemically defined medium with ribose as a sole carbon source (CDMR), *S. aureus* became a proline prototroph but remained as an arginine auxotroph (4). To confirm the results, we grew wild-type (WT) *S. aureus* strain Newman in CDM containing glucose (CDMG), ribose (CDMR), or no carbon source in the presence and absence of proline or arginine. As a control, the *ccpA* mutant was used. As shown in Fig. 1B, Δ*ccpA* grew in all conditions, whereas the WT strain failed to grow in CDMG in the absence of either proline or arginine, confirming that the proline and arginine synthesis pathways are under repression by CcpA. In the presence of ribose or without a carbon source, the WT strain grew without proline (-P in Fig. 1B), confirming that the proline auxotrophy of *S. aureus* is due to CCR. However, in the same conditions, the WT strain failed to grow without arginine (-R in Fig. 1B). In a previous study, *S. aureus* could grow without arginine when the growth medium contained sorbitol or arabinose as a sole carbon source. However, in our hands, *S. aureus* Newman still failed to grow in the conditions (-R in Fig. S1). Based on these results, we concluded that *S. aureus* could not synthesize arginine regardless of carbon sources.

### Arginine repressors are not involved in the arginine auxotrophic phenotype of *S. aureus*

In many bacteria, arginine repressors such as AhrC and ArgR repress arginine synthesis (12–16). *S. aureus* contains both AhrC and ArgR. To examine whether these arginine repressors are responsible for the *S. aureus* arginine auxotrophic phenotype in CDMR, we grew the WT strain and the mutants of the arginine repressors in CDMG or CDMR with or without arginine. As a control for CCR, the bacterial growth was also examined in the presence or absence of proline. In CDMR, all strains grew in the absence of proline; however, they failed to grow without arginine (-R in Fig. 2A), indicating that the arginine repressors are not responsible for the arginine auxotrophic phenotype of *S. aureus*. To identify the role of arginine repressors in arginine synthesis, we measured the transcript levels of the genes involved in arginine synthesis by qRT-PCR for WT and arginine repressor mutants. As shown, the transcription of *arcB* and *argG* was profoundly affected by the arginine repressor(s) (Fig. 2B). The transcription of *arcB* was dramatically increased only in the double mutant, indicating that the gene is redundantly repressed by AhrC and ArgR. Although the transcription of *argG* was increased in the *ahrC*-deletion mutant, it was more increased in the *argR* mutant, suggesting that *argG* is mainly repressed by ArgR. Transcription of all other genes was not largely affected by the arginine repressor mutations. Based on these results, we concluded that the arginine repressors control the conversion of ornithine to arginine but are not responsible for the arginine auxotrophy.

**Fig. 2.**
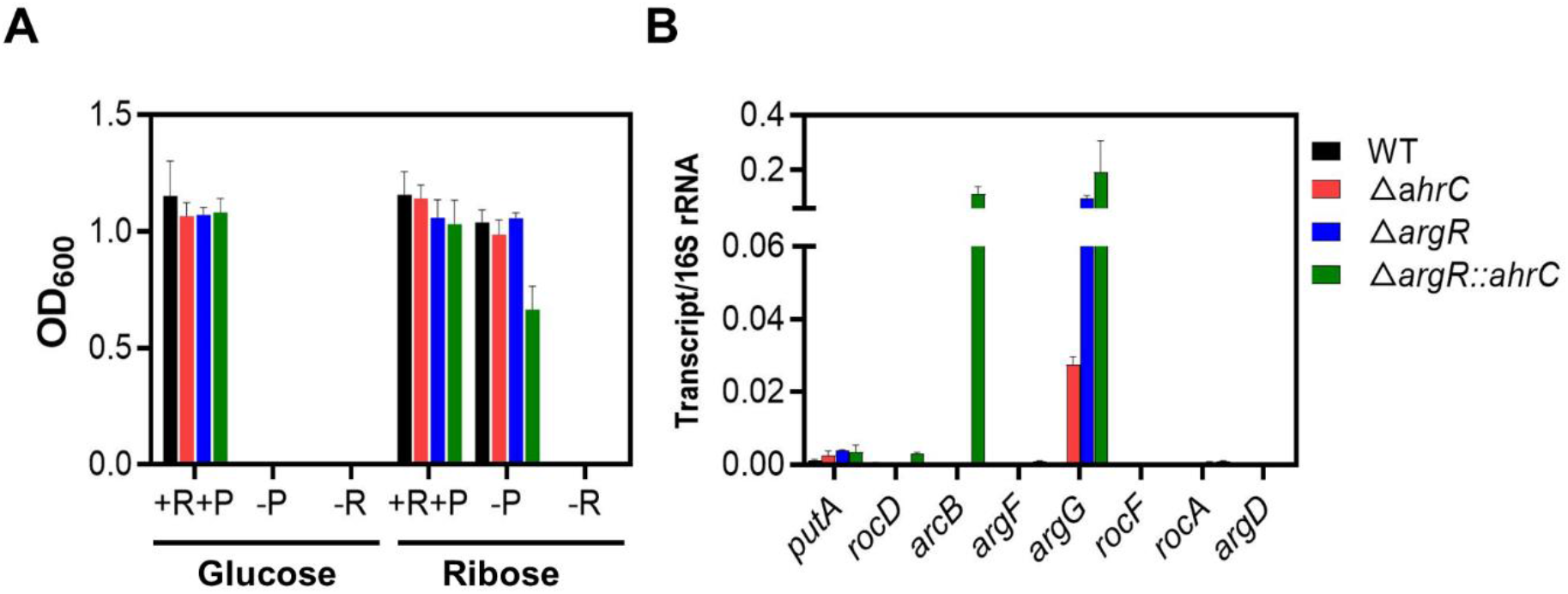
Arginine repressors do not control the arginine auxotrophy of *S. aureus*. (A) The effect of arginine repressor mutations on arginine and proline auxotrophy of *S. aureus. S. aureus* Newman wild-type (WT) and the arginine repressor mutants were grown at 37°C overnight in CDM containing glucose or ribose in the presence or absence of arginine and proline. +, presence; -, absence; R, arginine; P, proline. Data represent means ± SEM of three independent experiments. (B) The effect of arginine repressor mutations on the transcription of the genes involved in arginine metabolism. The test strains were grown in CDMG at 37°C until the exponential growth phase. The transcript levels of the test genes indicated under the graph were measured by qRT-PCR, where16S rRNA was used for internal control. The results were analyzed by the comparative C_T_ method. The experiments were performed in triplicate and repeated independently three times with similar results.

### In *S. aureus*, the proline-degradation step is responsible for the arginine auxotrophy

To determine which arginine synthesis steps are responsible for the arginine auxotrophy, we grew *S. aureus* Newman WT strain in CDMG, in which arginine was replaced by the arginine synthesis intermediate ornithine or citrulline. The WT strain grew in both growth media (Fig. 3A), indicating that staphylococcal arginine synthesis is impaired before the synthesis of ornithine (i.e., RocD or PutA).

**Fig. 3.**
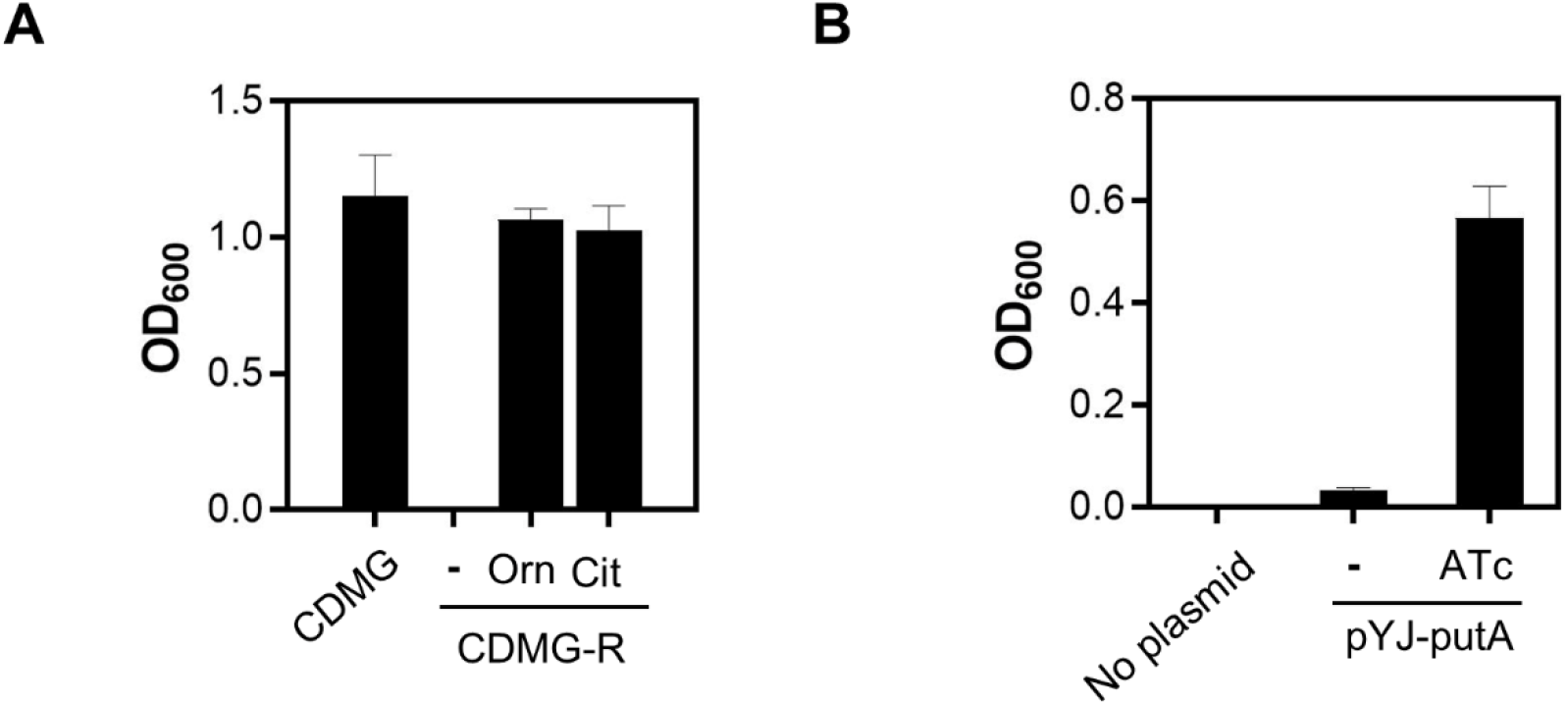
The proline-degradation step is responsible for the arginine auxotrophy of *S. aureus*. (A) Effect of arginine synthesis intermediates for staphylococcal growth without arginine. The Newman wild-type strain was grown at 37°C overnight in CDMG-R supplemented with ornithine (Orn) or citrulline (Cit). For a control, the strain was also grown in CDMG (i.e., CDM containing glucose and all amino acids). -, no addition. Data represent means ± SEM of three independent experiments. (B) Effect of the *putA* overexpression on arginine auxotrophy of *S. aureus*. *S. aureus* Newman containing the *putA* expression plasmid pYJ335-putA (pYJ-putA) was grown in CDMG-R in the absence (-) or presence of anhydrotetracycline (ATc). The Newman strain without the plasmid was used as a negative control. Data represent means ± SEM of three independent experiments.

The transcription of *rocD* and *putA* are both repressed by CcpA(3, 11). Of the two, RocD is shared by both arginine and proline synthesis pathways (Fig. 1A). Also, *S. aureus* could grow without proline when ribose was used as a sole carbon source (Fig. 1B), suggesting that the CCR imposed on *rocD* is released by ribose. Therefore, it is likely that the arginine auxotrophic phenotype in CDMR is due to the blockage at the proline degradation step by PutA. To test this hypothesis, we fused the *putA* gene to the anhydrotetracycline (ATc)-inducible promoter in a multi-copy plasmid pYJ335(17), and the resulting plasmid pYJ335-putA was inserted into the WT strain. In the absence of ATc, the strain barely grew; however, the addition of ATc significantly increased the growth of the bacterium (Fig. 3B), indicating that the insufficient production of PutA might be responsible for the *S. aureus* arginine auxotrophy in CDMR.

### In the wild-type strain, ribose appears to induce the expression of *putA* fully

In their previous study, Nuxoll et al. suggested that *S. aureus* arginine auxotrophy in CDMR is due to the inability of ribose to derepress CcpA (4). Indeed, the PutA production also seemed insufficient to support arginine synthesis in CDMR (Fig. 3B). To examine the possibility of the inadequate induction of *putA* in CDMR, we measured the *putA* transcripts levels in CDMG and CDMR for WT and the *ccpA*-deletion mutant. The *rocD* transcription was used as an indicator for CCR depression. In the WT strain, transcriptions of both *putA* and *rocD* were significantly increased in CDMR compared with CDMG (Fig. 4A), which was recapitulated in the promoter-lacZ fusion assay (Fig. 4B). These results demonstrate that, contrary to the previous postulation, the *putA* transcription is rather normally induced by ribose. In the *ccpA* mutant, both *putA* and *rocD* showed significant transcription in CDMG (Fig. 4A), further confirming that both genes are repressed by CcpA. Importantly, the *putA* transcript level in CDMR was slightly higher than that in CDMG of *ccpA* mutant (Fig. 4A). Considering the fact that the *ccpA* mutant grows in CDMG without arginine, these results indicate that inadequate induction of *putA* might not responsible for the *S. aureus* arginine auxotrophy. This conjecture was further supported by Western blot analysis for PutA. The PutA level of WT in CDMR was similar to or slightly higher than that of the *ccpA* mutant in CDMG (Fig. 4C). To further investigate the *putA* induction in CDMR, we generated a single copy *putA* complementation plasmid pROKA-putA. Then the CRE-sequence or the entire promoter of *putA* was replaced with those of *rocD*. The original and variant plasmids were inserted into WT, Δ*putA*, and Δ*ccpA* strains, and the resulting strains were grown in CDMR with or without arginine (CDMR-R). As shown, regardless of the complementary plasmids, the WT strains failed to grow in the absence of arginine (Fig. 4D). Based on these results, we concluded that *putA* seems to be fully induced in CDMR, and inadequate induction of *putA* is not responsible for *S. aureus* arginine auxotrophy.

**Fig. 4.**
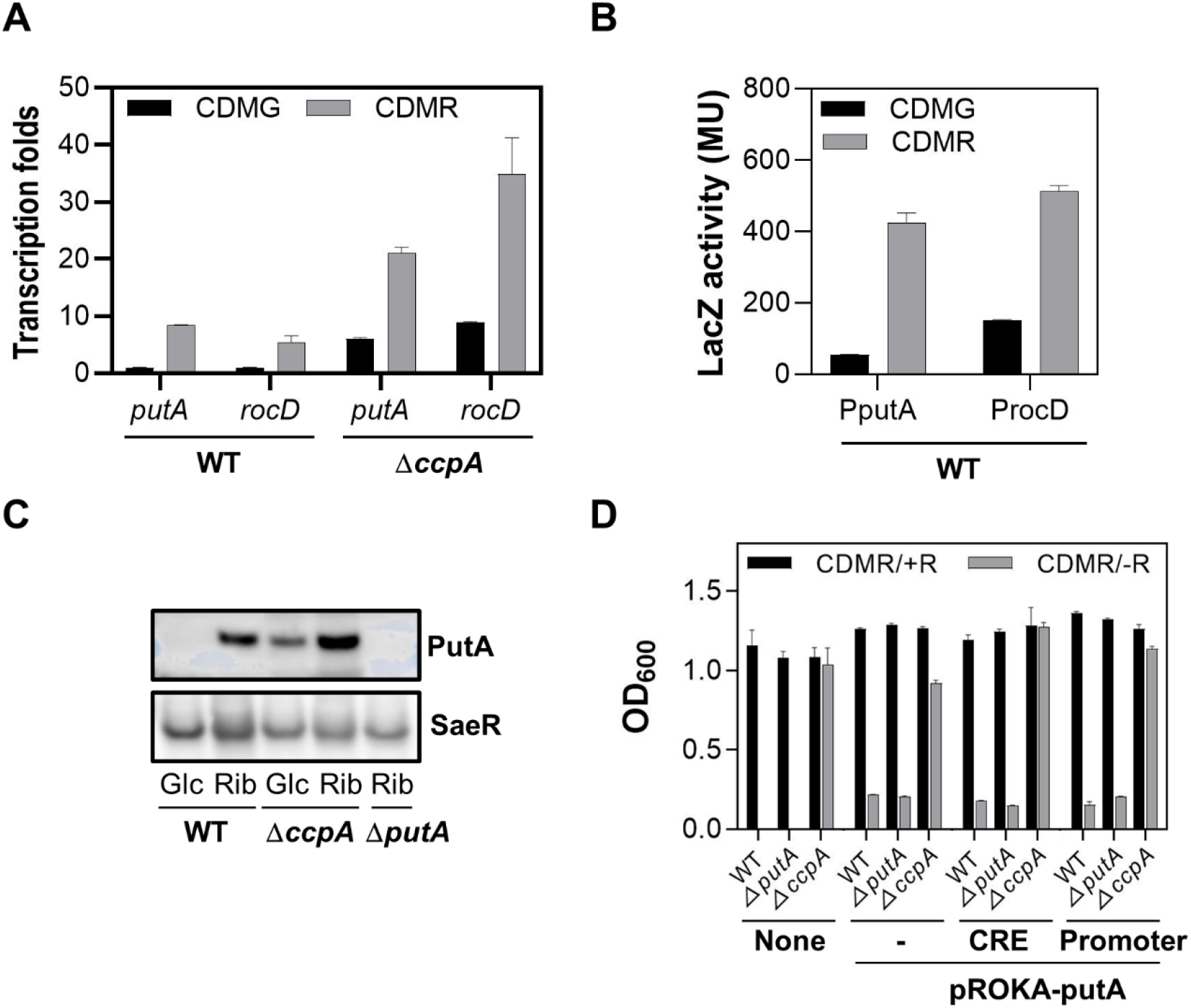
Ribose fully induces the expression of PutA. (A) Ribose-mediated induction of *putA* and *rocD* transcription measured by qRT-PCR analysis. The wild-type (WT) and the *ccpA* mutant (Δ*ccpA*) were grown in CDMG or CDMR until the exponential growth phase, and total RNA was purified. 16S rRNA was used as an internal control, and the results were analyzed by the comparative C_T_ method and normalized to the WT value in CDMG. (B) The ribose-mediated induction of the *putA* transcription measured by the promoter-lacZ fusion assay. Newman WT strain containing the *putA* promoter-*lacZ or* the *rocD* promoter-*lacZ* plasmid was grown in CDMG or CDMR until the exponential growth phase. Cells were collected and lysed; then, LacZ activity was measured. All assays were performed in triplicate and repeated independently three times. (C) Carbon source effect on the expression of the PutA protein. WT and Δ*ccpA* strain were grown in CDMG or CDMR, and harvested at the exponential growth phase. The expression of PutA was measured by Western blotting with a PutA antibody. A *putA* transposon mutant strain (Δ*putA*) was used for negative control. For loading control, SaeR, the response regulator of the SaeRS two-component system, was detected. Glc, CDMG; Rib, CDMR. (D) Effect of the CRE sequence or the promoter replacement on arginine auxotrophy of *S. aureus*. The CRE sequence or the entire promoter sequence of the *putA* promoter in pROKA-putA was replaced with that of the *rocD* promoter. The resulting plasmids were inserted into the wild-type (WT), the *putA*-deletion mutant (Δ*putA*), and the *ccpA*-deletion mutant (Δ*ccpA*). The test strains were grown at 37°C overnight in CDMR and CDMR-R. -, no change; CRE, the CRE sequence replacement; promoter, the promoter replacement. Data represent means ± SEM of three independent experiments.

### Uncharacterized repeat sequences are involved in the ribose-mediated activation of the *putA* promoter

In the previous experiment, we observed that both *putA* and *rocD* transcript levels in Δ*ccpA* were further increased in CDMR compared with CDMG (see Δ*ccpA* in Fig. 4A), indicating that transcription of those genes was activated by ribose in a CCR-independent manner. The *putA* promoter and its upstream region contain three types of *cis*-elements: two ARG boxes (i.e., the binding sequence for arginine repressors), uncharacterized two identical repeat sequences (RS1 and RS2), and a CRE sequence (Fig. 5A). One of the ARG boxes overlaps with −10 sequence of the putative promoter for NWMN_1657, whereas the CRE sequence is right next to one of the repeat sequences and overlaps with −35 sequence of the *putA* promoter. To investigate which of those *cis*-elements is responsible for the CCR-independent regulation, we generated various deletions of the *cis*-elements in the *putA*-lacZ fusion plasmid (Fig. 5B). Those plasmids were inserted into the Newman WT strain, and the LacZ activity was measured in CDMG or CDMR. The deletion of the ARG boxes did not significantly affect the LacZ activity in both growth media, showing that the ARG boxes do not affect the *putA* promoter activity. On the other hand, the deletion of the uncharacterized repeat sequences additively lowered the LacZ activity in CDMR (Δ4 and Δ5 in Fig. 5B and 5C). As expected, when half of the CRE-sequence was deleted without affecting the −35 sequence, the LacZ activity was increased in CDMG but not in CDMR (Δ6 in Fig. 5B and 5C). The *putA* core promoter (i.e., the promoter in the Δ6 mutant) showed only slightly higher LacZ activity in CDMR than in CDMG. Based on these results, we concluded that the uncharacterized repeat sequences are responsible for the ribose-mediated activation of the *putA* promoter.

**Fig. 5.**
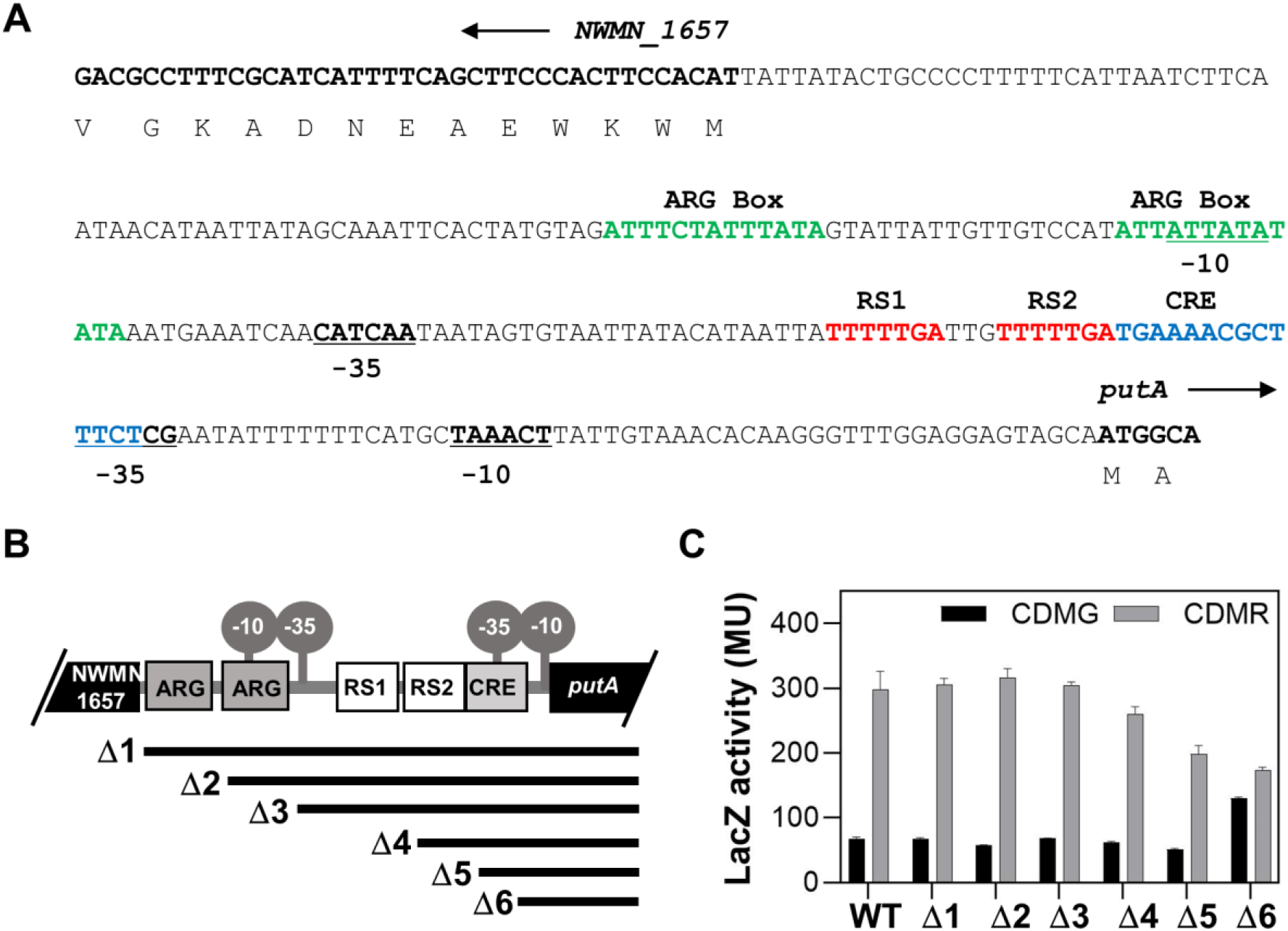
Uncharacterized repeat sequences are involved in the ribose-mediated activation of the *putA* promoter. (A) The DNA sequence of the *putA* promoter region. ARG Box, the putative binding site of arginine repressors; RS, repeat sequence. The promoter sequences (−10 and −35) are also indicated. The *putA* promoter sequence was verified by mutagenesis analysis, whereas the promoter sequence for NWMN-1657 was predicted. (B) The *cis*-element deletions in pOS1-PputA-lacZ. The remaining putA promoter sequences are indicated as a thick line for each deletion mutant. (C) The effect of *cis*-element deletion on the *putA* promoter activity. *S. aureus* Newman carrying one of the pOS1-PputA-lacZ plasmids was grown at 37°C in CDMG or CDMR, and LacZ activity was measured at the exponential growth phase. WT, no deletion in pOS1-PputA-lacZ; Δ1 - Δ6, *cis*-element deletion mutants of pOS1-PputA-lacZ shown in (B). Data represent means ± SEM of three independent experiments.

### The intracellular concentration of pyrroline-5-carboxylate (P5C) is the determining factor for arginine synthesis

So far, the experimental results indicate that, although the transcription of *putA* is fully induced by ribose, *S. aureus* remains arginine auxotroph in CDMR. To find the reason for the arginine auxotrophy, we decided to identify arginine prototrophic mutants. The mix of 1,736 transposon mutants of *S. aureus* Newman (i.e., Phoenix library) was spread on the CDMR agar plate without arginine (CDMR-R agar) and incubated for two days (18). The colonies on the plate were then inoculated in CDMR-R broth to confirm their arginine prototrophy. Inverse PCR analysis of those mutants found that they have a transposon insertion in *proC*, the gene encoding pyrroline-5-carboxylate (P5C) reductase. This enzyme converts P5C into proline, the last step of the proline synthesis and the reverse reaction of proline degradation (Fig. 1A). The *proC* mutant grew in CDMR-R and CDMG-R, indicating that arginine synthesis in the strain is independent of CCR (Fig. 6A). When the transposon mutation was transduced into the strain USA300, a predominant MRSA strain in the US, the resulting mutant could grow without arginine, indicating that the *proC* mutation converts *S. aureus* into arginine prototroph regardless of the strain background (Fig. S2).

**Fig. 6.**
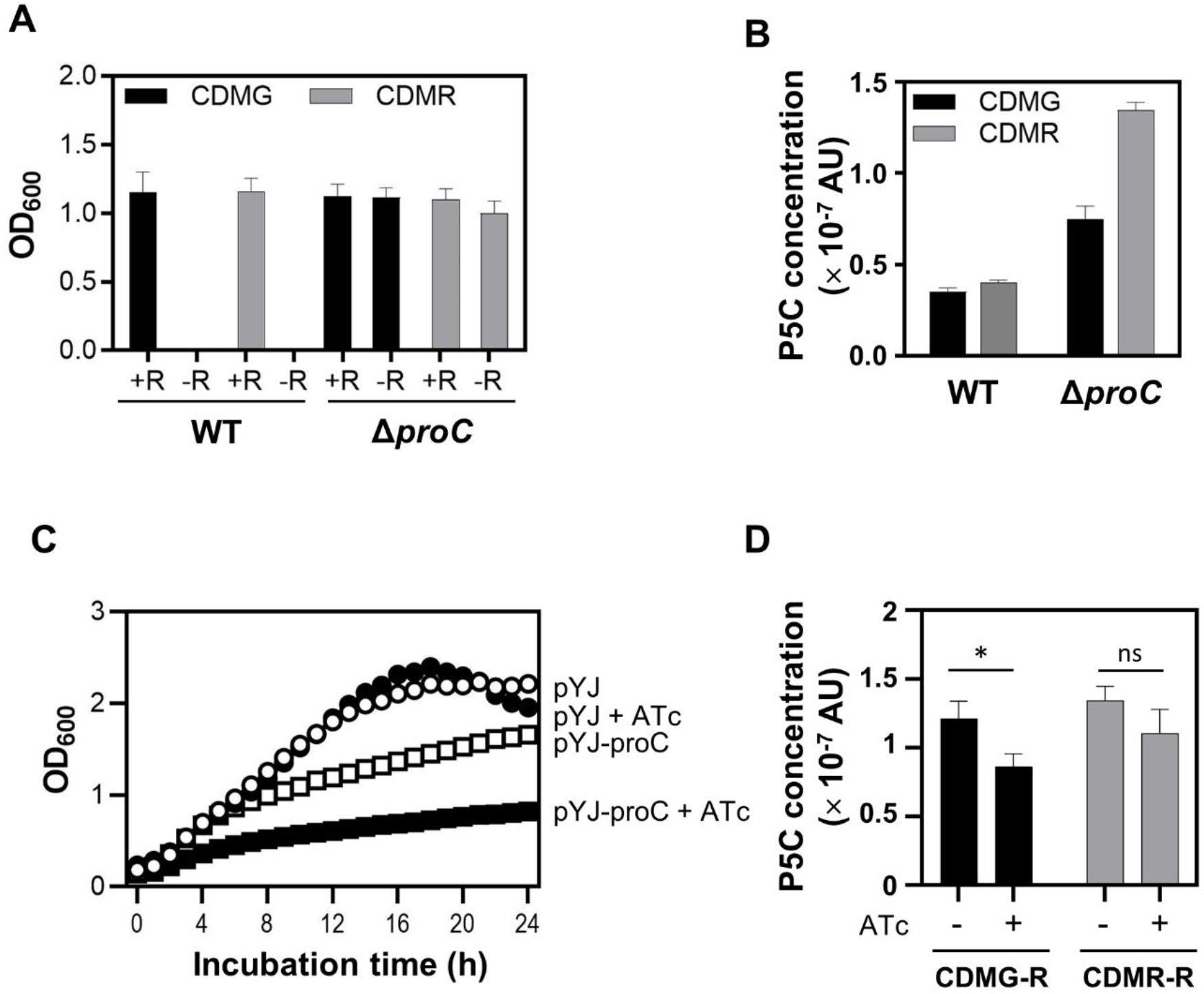
The pyrroline-5-carboxylate (P5C) intracellular concentration is the determining factor for arginine synthesis. (A) Effect of the *proC* disruption on the arginine auxotrophy of *S. aureus*. *S. aureus* Neman wild-type (WT) and *proC*-deletion mutant (Δ*proC*) were grown at 37°C overnight in CDMG or CDMR with or without arginine. + R, presence of arginine; - R, no arginine. Data represent means ± SEM of three independent experiments. (B) The intracellular concentration of pyrroline-5-carboxylate (P5C) in WT and Δ*proC*. Cells were grown at 37°C overnight in CDMG and CDMR; then, the concentration of P5C was measured as described in Methods. AU, arbitrary unit. (C) Effect of *proC* overexpression on the growth of the *ccpA*-deletion mutant (Δ*ccpA*) in CDMR-R (i.e., CDMR without arginine). The Δ*ccpA* strain carrying the *proC*-expression plasmid pYJ335-proC was grown at 37°C for 24 h. For clarity, the error bars are not shown. The growth curves of Δ*ccpA* in CDMG-R is provided in Fig. S4. pYJ, pYJ335; pYJ-proC, pYJ335-proC; + ATc, addition of anhydrotetracycline (100 ng/ml). (D) Effect of *proC* overexpression on the intracellular P5C concentration of Δ*ccpA*. The Δ*ccpA* strain carrying the *proC*-expression plasmid pYJ335-proC was grown at 37°C overnight. ATc, anhydrotetracycline; -, absence; +, presence. Data represent means ± SEM of three independent experiments. Statistical significance was assessed by unpaired, two-tailed Student’s t-test. *, p < 0.05; ns, not significant.

A straightforward explanation for the arginine prototrophy of the *proC* mutant is that the disruption of *proC* increases the intracellular concentration of P5C, mimicking the overexpression of PutA (Fig. 3B). To examine this hypothesis, we measured the intracellular levels of P5C in WT and Δ*proC* using the ninhydrin derivatization method (19, 20). Indeed, the concentration of P5C was significantly increased in the *proC* transposon mutant, regardless of the carbon source (Fig. 6B). When the transcription of *putA* was induced from the multi-copy plasmid pYJ335-putA, which converted *S. aureus* into arginine prototroph (Fig. 3B), the intracellular concentration of P5C was also increased (Fig. S3).

To further test whether the intracellular concentration of P5C is a critical determinant for the arginine prototrophy, we generated pYJ335-proC, where *proC* is transcribed from an ATc-inducible promoter. The plasmid was inserted into Δ*ccpA*, which can grow without arginine, and the resulting strain was grown in CDMG-R and CDMR-R. As shown, the inducer alone did not significantly affect the growth of Δ*ccpA* (pYJ vs. pYJ+ATc in Fig. 6C and Fig. S4). However, when *proC* transcription was induced by ATc, the growth of Δ*ccpA* was significantly inhibited (pYJ-proC + ATc in Fig. 6C and Fig. S4). When the intracellular P5C concentration was measured, it was lower in the ProC-overexpression strains, although the difference was not statistically significant in CDMR-R due to the high variability of the measurements (Fig. 6D). Based on these results, we concluded that the intracellular concentration of P5C is the determining factor for arginine prototrophy.

### The *argDCJB* operon is functional if transcribed

The *argDCJB* operon encodes enzymes for arginine synthesis from glutamate (Fig. 1A). However, the operon is known not to be transcribed (4, 10). To examine whether the operon is functional if transcribed, we fused the entire *argDCJB* operon to the ATc-inducible promoter in pYJ335, and inserted the resulting plasmid into the WT *S. aureus* Newman. In CDMG-R, the strain grew when the inducer ATc was added (Fig. 7), demonstrating that, once transcribed, those genes can produce functional enzymes that confer arginine prototrophy to *S. aureus*.

**Fig. 7.**
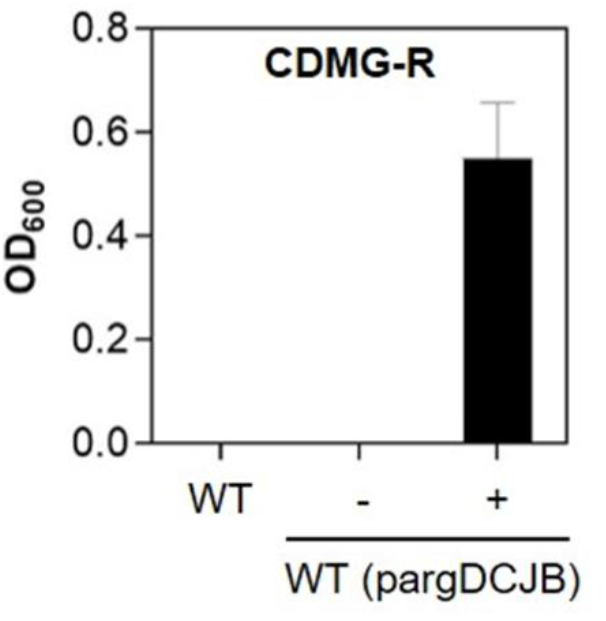
The *argDCJB* operon is functional once transcribed. *S. aureus* Newman containing pYJ335-argDCJB was grown in CDMG-R with (+) or without (-) anhydrotetracycline (100 ng/ml). For negative control, wild-type (WT) Newman strain without the plasmid was used. pargDCJB, pYJ335-argDCJB. Data represent means ± SEM of three independent experiments.

### The arginine/proline’s role as alternative carbon sources does not contribute to the *in vivo* survival of *S. aureus*

In *S. aureus*, arginine and proline can be fed into the TCA cycle through glutamate in the absence of glucose (Fig. 1A)(11). Next, we decided to examine whether the role of these amino acids as alternative carbon sources is critical for staphylococcal virulence and survival during infection. To block the synthesis of P5C, we disrupted both *rocD* and *putA*. We also deleted *rocA* to stop the flow from P5C to glutamate. These two mutant strains were administered into mice via a retro-orbital route with the WT strain. At 3 d post-infection, the bacterial loads in the kidney, liver, and heart were measured. As shown in Fig. 8, none of the mutations reduced the bacterial loads in the organs. On the contrary, the CFU of the Δ*rocD::putA* double mutant was slightly but significantly higher than that of WT in the kidney. These results demonstrate that those amino acids’ role as alternative carbon sources is, at least, not essential for the virulence and survival of *S. aureus*.

**Fig. 8.**
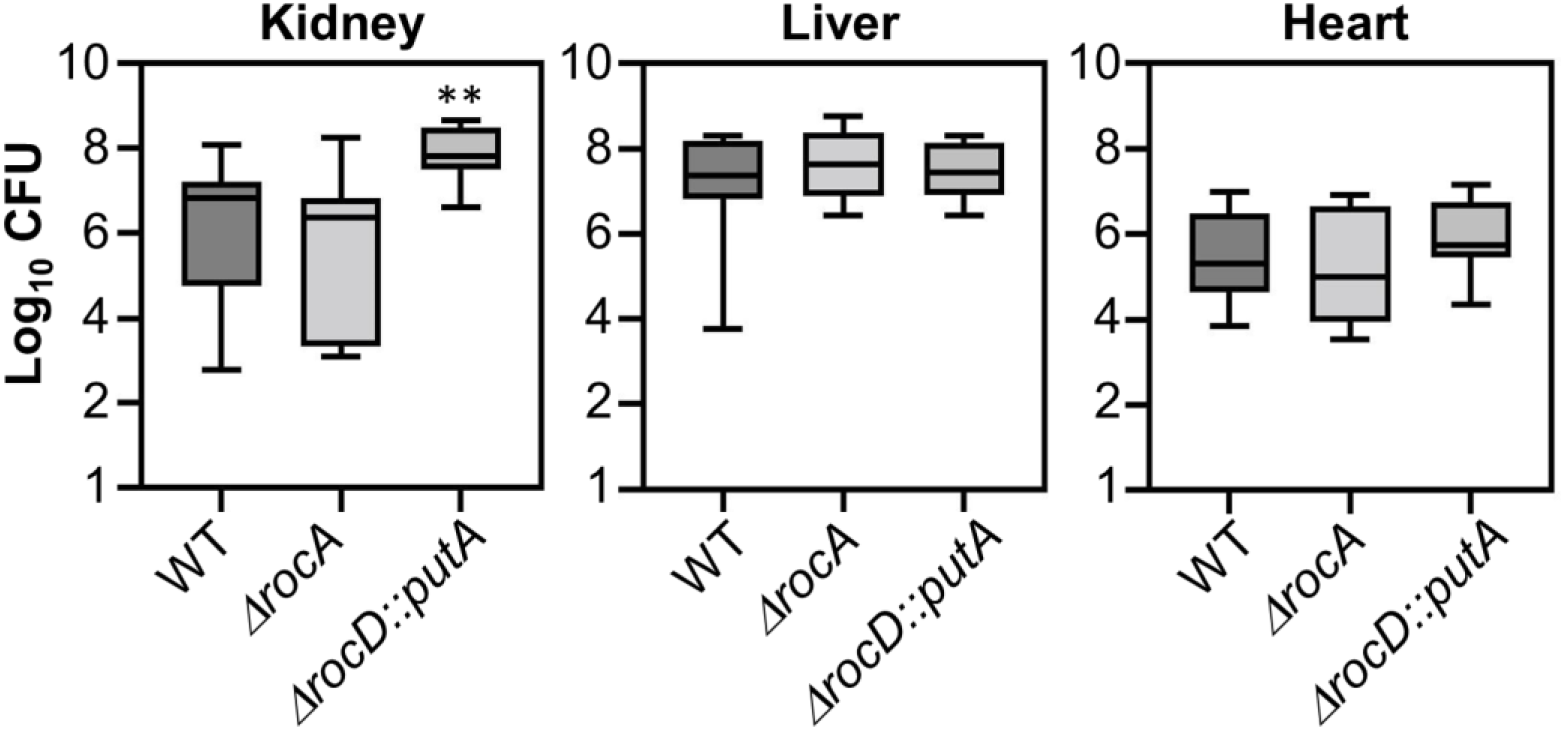
The arginine-proline interconversion pathway does not contribute to the virulence and survival of *S. aureus*. The test strains (10^7^ CFU) were injected into 6 – 12 C57BL/6 mice via retro-orbital route. At 3 days post-infection, the bacterial loads in the organs were measured by CFU counting. The CFU in the kidney and liver were counted from 12 mice, whereas the heart CFU counting was from six mice. Statistical significance was assessed by unpaired, two-tailed Student’s t-test. **, p < 0.005.

## Discussion

The genome of *S. aureus* harbors genes for the synthesis of all 20 amino acids (21). However, due to CCR by CcpA, *S. aureus* is phenotypically auxotrophic to proline (3). *S. aureus* is also known to be auxotrophic to arginine due to the CcpA-mediated CCR (4). Indeed, the *ccpA* mutant does not need arginine for growth (Fig. 1B) (4, 11). However, the disruption of CCR with non-preferred carbon sources failed to convert *S. aureus* into arginine prototroph (Fig. 1B)(11). In this study, we demonstrated that the intracellular concentration of P5C, not CCR, is the determining factor for arginine auxotrophy, and the reduction of P5C concentration by the P5C reductase ProC is responsible for the *S. aureus* arginine auxotrophy. Therefore, it is unlikely that *S. aureus* can synthesize arginine from proline under physiological conditions without artificial gene knockouts.

To explain *S. aureus* arginine auxotrophy in CDMR, Nuxoll et al. suggested that ribose is unable to derepress CcpA (4). Initially, we also hypothesized that the inadequate induction of the *putA* gene by ribose is responsible for the arginine auxotrophy in CDMR. However, the transcription of *putA* seems to be fully induced by ribose. The *putA* transcription was increased eight folds in CDMR compared with CDMG, while the *rocD* transcription was increased 3 – 6 folds (Fig. 4A and 4B). Also, the transcription from the *putA* promoter was not significantly affected by the deletion of the CRE sequence, the CcpA binding site, when the bacterium was grown in CDMR (Δ5 vs. Δ6 in Fig. 5C). Therefore, it is likely that ribose fully derepresses CcpA, and CCR is not responsible for the arginine auxotrophy.

Deletion mutagenesis showed that the *putA* promoter is regulated differently depending on the carbon source. In CDMG, the activity of *putA* promoter was solely controlled by CcpA (Δ5 vs. Δ6 in Fig. 5C), whereas, in CDMR, it was regulated by two repeat sequences (RS1 and RS2 in Fig. 5A and Δ3 - Δ5 in Fig. 5C). Those repeat sequences might be binding sites for a hitherto unknown transcription factor. Since the repeat sequences are right next to CRE, the DNA bindings of the putative transcription factor and CcpA are likely to be mutually exclusive. Indeed, in CDMG, the deletion of the repeat sequence did not affect the activity of the *putA* promoter, whereas, in CDMR, the deletion of CRE showed little effect on the *putA* promoter activity (Fig. 5C). It should also be noted that the distance between RS2 and the −35 sequence of the *putA* promoter is 10 bp, a nucleotide number for one helix turn. Therefore, once bound to DNA, the putative transcription activator will be on the same side as RNA polymerase, a prerequisite for direct interaction. Intriguingly, each repeat sequence contributed to the *putA* promoter activity in an additive manner (Δ4 and Δ5 in Fig. 5C). Therefore, it is possible that the putative transcription factor binds to the sequence as a monomer. However, they are just speculations at this stage, and more studies are required to confirm the existence of such a transcription factor.

In *S. aureus*, the P5C concentration appears to be the key determinant for arginine synthesis from proline. The increased P5C concentration in Δ*proC* allowed the strain to grow without arginine regardless of the carbon sources, whereas the decreased P5C concentration inhibited the arginine prototrophy of the *ccpA* mutant (Fig. 6 and Fig. S4). Since RocA also uses P5C as a substrate to synthesize glutamate, the Δ*rocA* mutant is expected to have an increased concentration of P5C. Indeed, in CDMR, the P5C concentration in Δ*rocA* was higher than that in WT but lower than that in either Δ*ccpA* or Δ*proC* mutant (Fig. S5A). However, the Δ*rocA* mutant barely grew in CDMR without arginine (Fig. S5B), suggesting that the P5C concentration in Δ*rocA* is not sufficient to support arginine synthesis (Fig. 1A). Intriguingly, when *S. aureus* was supplemented with ^13^C-labeled proline in CDM without carbon source, it produced ^13^C-labeled ornithine but not citrulline (11), indicating that, even at the concentration insufficient to support arginine synthesis, P5C still can be converted to ornithine by RocD. However, the concentration of the produced ornithine is likely too low to be further processed by ArgG into citrulline (Fig. 1A).

Because of the arginine prototrophy of the *ccpA* mutant, *S. aureus* has been known to synthesize arginine from proline via the CCR control (4). However, when the CcpA-mediated CCR is abolished by secondary carbon sources, *S. aureus* still remains arginine auxotroph (Fig. 1B and Fig. S1). Also, like the *ccpA* deletion, the knockout mutation of *proC* converts *S. aureus* into arginine prototroph (Fig. 6A and 6B). Therefore, the arginine prototrophic phenotype of the *ccpA* mutant is more an artifact of the gene disruption than a regulatory outcome of the derepression of CCR. As Halsey et al. clearly showed, it is likely that the proline catabolite pathway functions to provide an alternative carbon source in the absence of glucose, not to synthesize arginine (11).

Commonly, bacteria synthesize arginine from glutamate using the *argDCJB* operon. Although the *S. aureus* genome contains the *argDCJB* operon, the operon appears to be dormant. Mader et al. has reported that the *argDCJB* operon was not transcribed in *S. aureus* HG001, a derivative of NCTC 8325, under 44 different growth conditions (10). However, when the operon was transcribed from an ATc-inducible promoter, *S. aureus* became an arginine prototroph (Fig. 7), indicating that the genes in the operon are intact. Without significant contribution to bacterial growth and survival, it is unlikely that the entire operon survives natural selection and remains intact. Therefore, it is reasonable to predict that *S. aureus* requires the *argDCJB* operon to synthesize arginine under certain environmental conditions. In their study, Mader et al. identified 312 genes never expressed in 44 different growth conditions (10). Intriguingly, those genes include *arcB*, whose transcription is repressed by ArgR/AhrC (Fig. 2B), and *vraDE*, whose transcription is induced by bacitracin and nisin via the BraSR two-component system (22). Therefore, those 44 different growth conditions are not comprehensive and did not include the growth condition where the transcription of the *argDCJB* operon is induced. Considering that, without mutation in *ccpA* or *proC*, the P5C concentration is too low for arginine synthesis (Fig. 6B), ArgDCJB likely constitutes the genuine pathway for arginine synthesis in *S. aureus*.

Finally, we propose the following models for the effect of CCR and arginine on the arginine-proline interconversion pathway (Fig. 9). In CDMG, almost all steps are repressed either by the CcpA-mediated CCR or by the arginine repressors. Although the conversion of P5C to proline by ProC is not repressed, proline is not synthesized due to the low concentration of P5C. Therefore, *S. aureus* is auxotrophic to arginine and proline in this condition. In CDMR, CCR is abolished, but due to the presence of arginine, the arginine repressors are functional and repress the conversion of ornithine to arginine. In this condition, both arginine and proline are converted into P5C and fuel the TCA cycle through glutamate synthesis by RocA. Due to the constitutive activity of ProC, P5C is also converted to proline, leading to proline prototrophy. In CDMR-R (i.e., CDMR without arginine), neither CcpA nor arginine repressors are function, and all steps are open. Although proline can be converted to P5C by PutA and might be further processed to glutamate, due to the constant consumption of P5C by ProC, the intracellular P5C concentration is too low to produce sufficient amount of ornithine for arginine synthesis, and *S. aureus* fails to grow. In summary, *S. aureus* cannot synthesize arginine from proline in any of these three conditions.

**Fig. 9.**
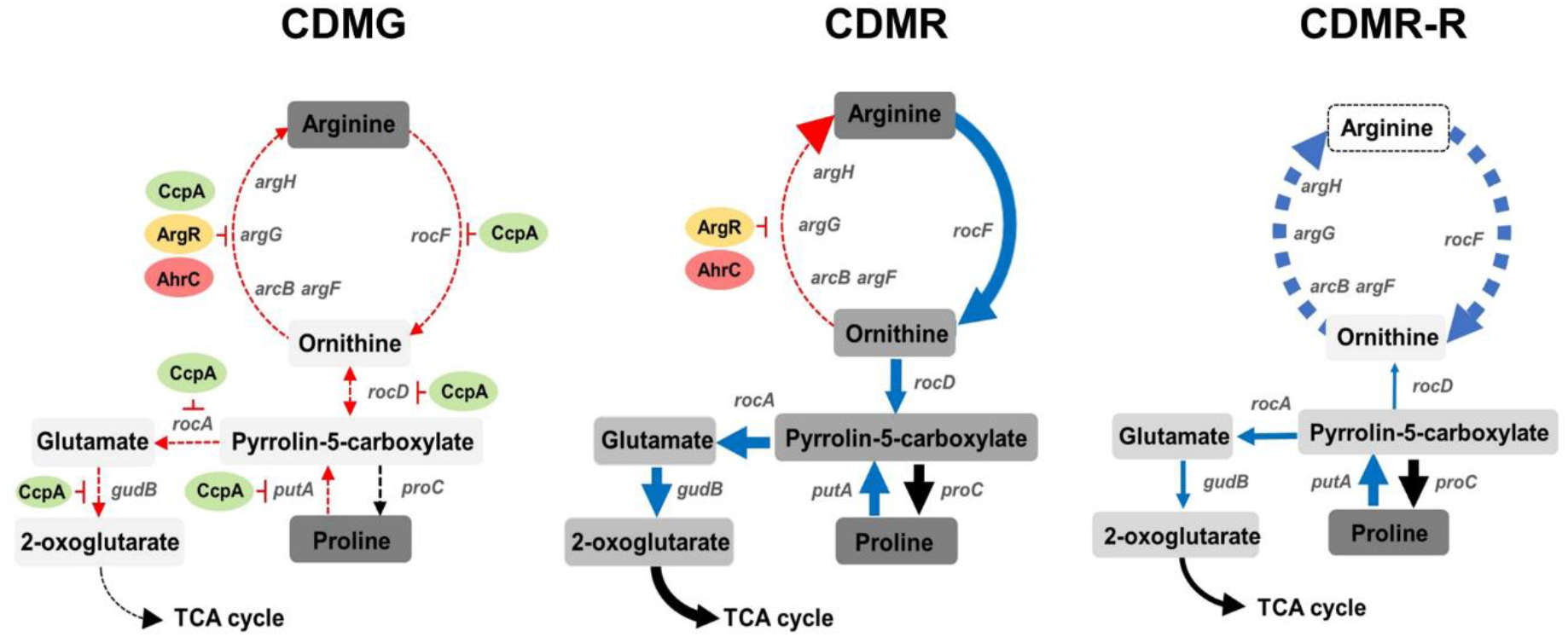
Models for the regulation of arginine and proline metabolism by carbon catabolite repression and arginine repressors. Red arrow, repressed; blue arrow, induced; black arrow, constitutive. The dashed lines indicate non-functional steps. The gradient of gray colors represents relative concentrations.

## Materials and Methods

### Ethics Statement

All animal experiments were performed in accordance with the Guide for the Care and Use of Laboratory Animals of the National Institutes of Health. The animal protocol was approved by IUSM-NW IACUC (protocol # NW-48). Every effort was made to minimize the suffering of the animals.

### Bacterial strains and growth conditions

Bacterial strains used in this study are listed in Table 1. For bacterial culture in chemically defined medium (CDM)(23, 24), single colonies on tryptic soy agar (TSA) were inoculated in tryptic soy broth (TSB), and the culture was incubated at *37°C* with shaking overnight. Cells were collected by centrifugation, washed with sterile PBS, and suspended in sterile water. The cell suspension (1% of the culture volume) was inoculated into CDM and incubated at 37°C with shaking. For transduction of transposon mutation, heart infusion broth (HIB, BD Difco) supplemented with 5 mM CaCl_2_ (Sigma) was used. *Escherichia coli* was used for genetic manipulation of plasmids and grown in lysogeny broth (LB, BD Difco). When necessary, antibiotics were added to the cultures at the following concentrations: ampicillin, 100 μg/ml; chloramphenicol, 15 μg/ml; erythromycin, 10 μg/ml.

**Table 1.**
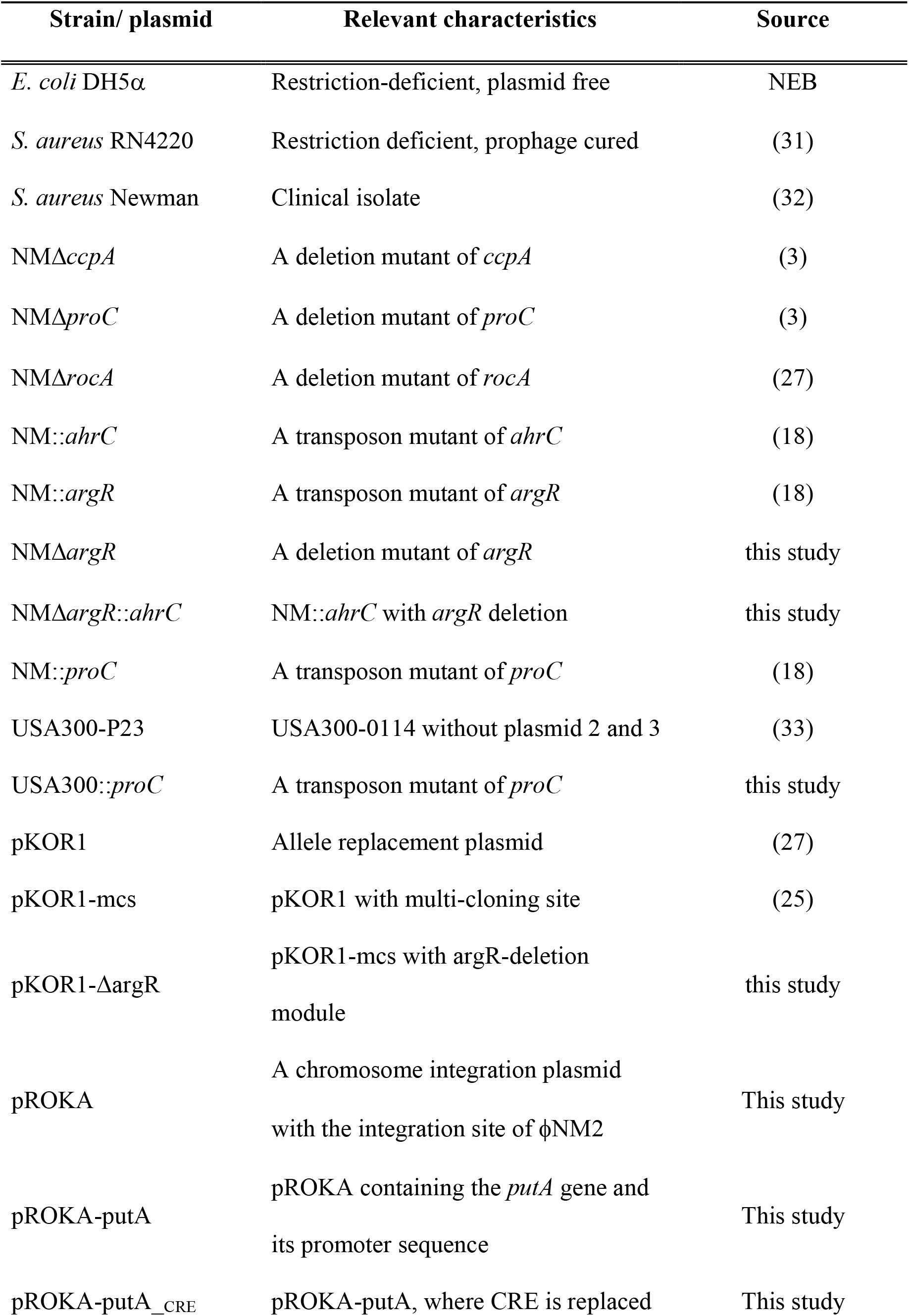
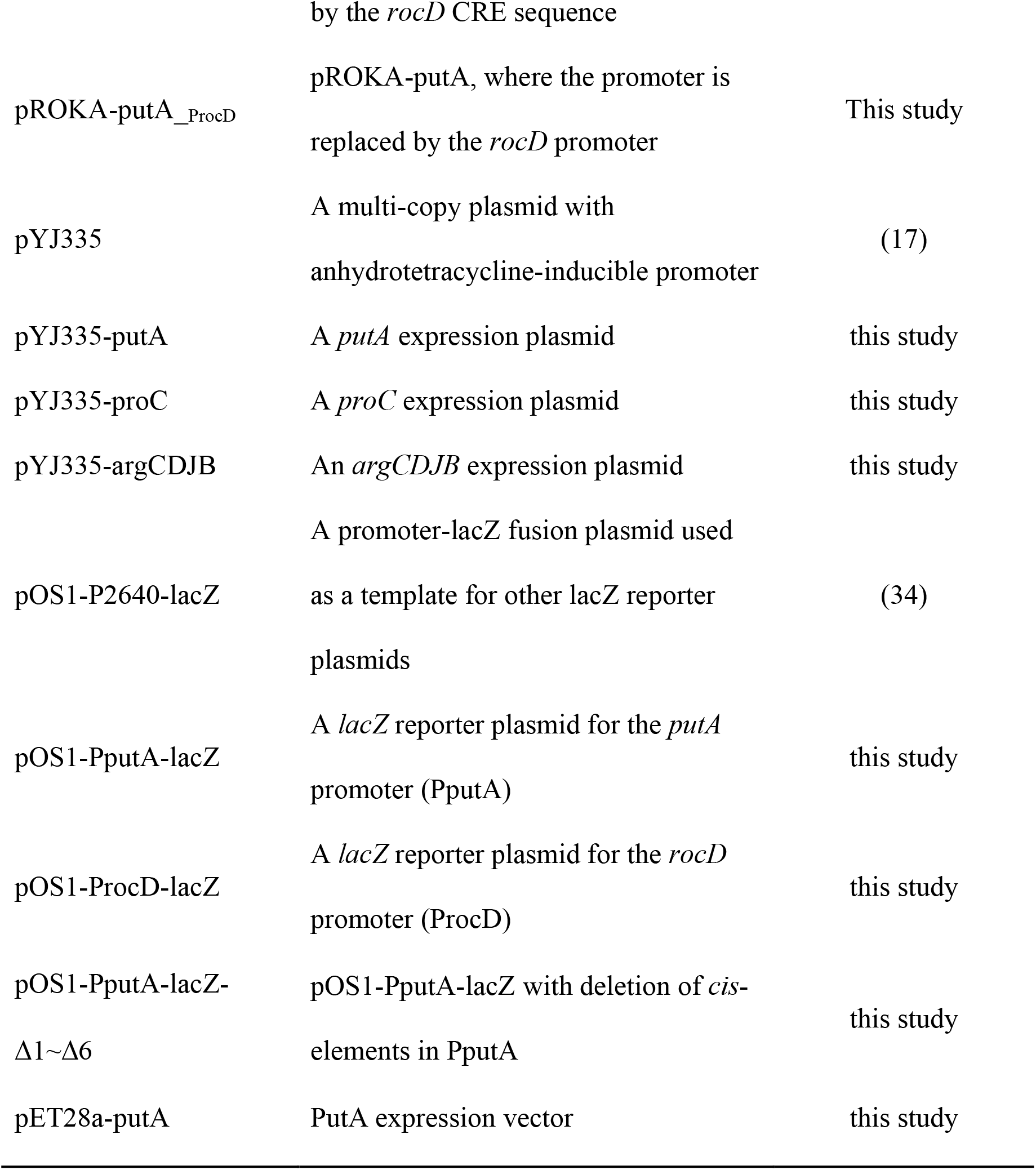
Bacterial strains and plasmids used in this study.

### Generation of an *ahrC/argR* double mutant

To delete *argR*, we used the plasmid pKOR1-mcs (25). 1 kb flanking sequences of *argR* were PCR-amplified with the primer sets P1742/P1743 and P1744/P1755 (Table 2), whereas pKOR1-mcs was amplified with the primer set P1740/P1741 (Table 2). The PCR products were purified and joined with ligation-independent cloning (26)and inserted into *E. coli* DH5α by transformation. After confirmation of the DNA sequences of the inserted DNA by DNA sequencing, the resulting plasmid, pKOR1-ΔargR was inserted into *S. aureus* RN4220 by electroporation and subsequently into *S. aureus* Newman by transduction with ϕ85. Deletion of *argR* was carried out as described previously(27). Then, the transposon in NM::*ahrC* was transduced into the *argR* deletion mutant, resulting in NMΔ*argR::ahrC* (Table 1).

**Table 2.**
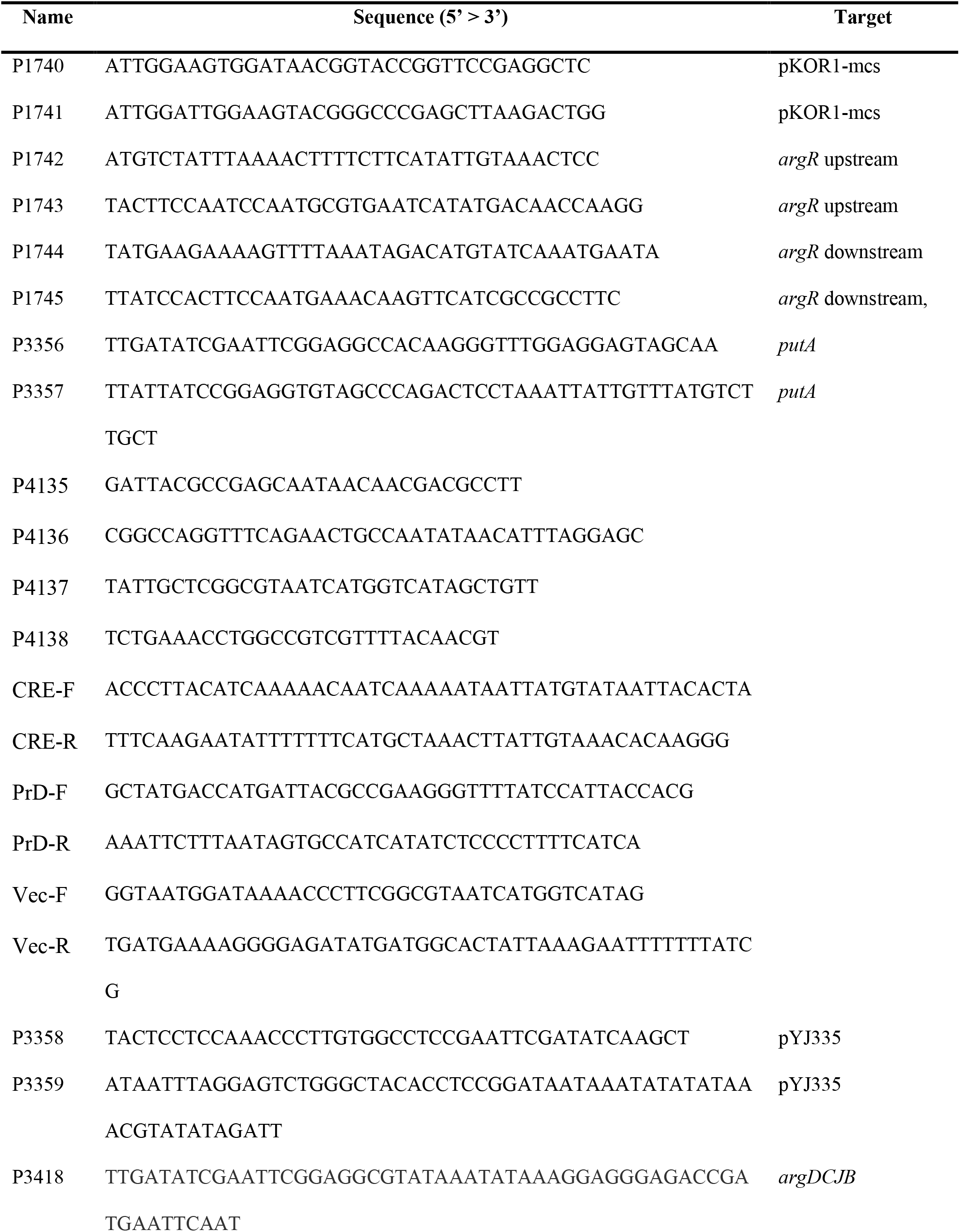
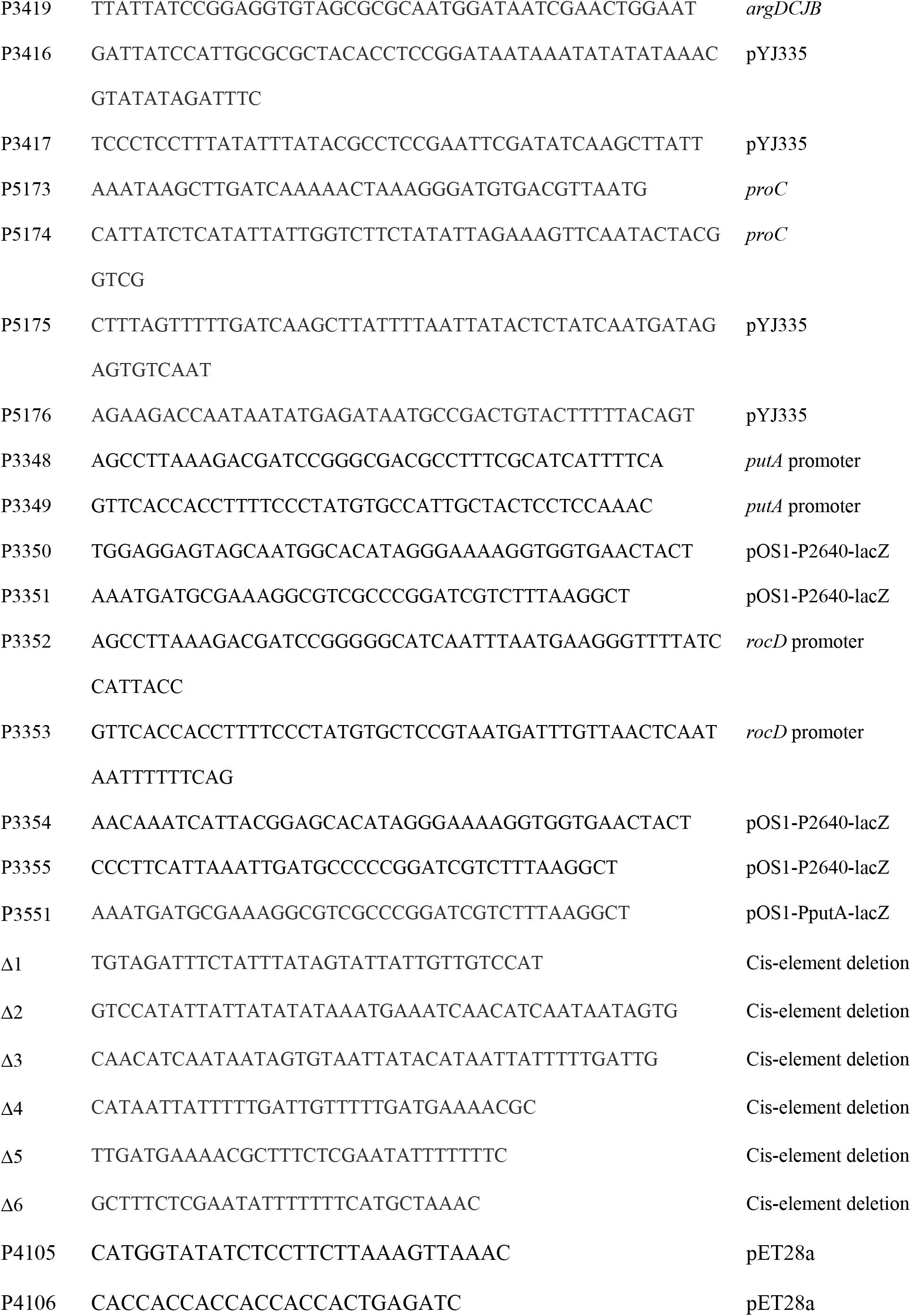
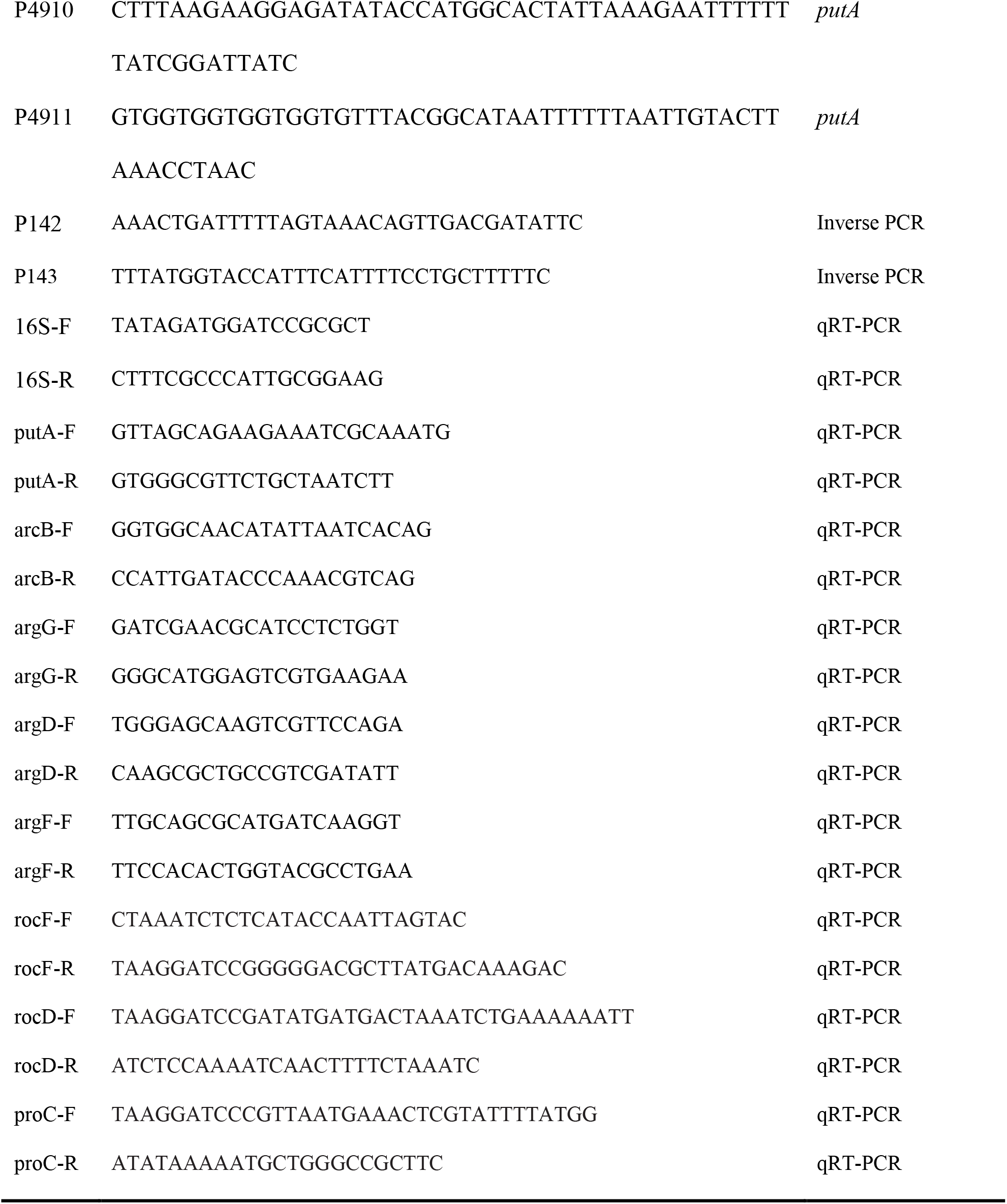
Oligonucleotides used in this study.

### Construction of gene expression plasmids

pROKA is a chromosome integration plasmid containing p15A origin for the replication in *E. coli* and the integrase gene and phage attachment sequence (attP) of ϕNM2 in *S. aureus* Newman. The plasmid integrates into the intergenic region between *rpmF* and *isdB* (28).

To generate pROKA-putA, we PCR-amplified the *putA* gene with primer pair P4135/P4136 (Table 2). pROKA was also PCR-amplified with primer pair P4137/P4138 (Table 2). The PCR products were treated with *Dpn*I (NEB) at 37°C for 30 minutes, purified with a PCR purification kit (Qiagen), and assembled by the Gibson method (29). The assembled plasmid was inserted into *E. coli* DH5α by transformation.

To replace the CRE sequence in pROKA-putA with that of *rocD*, we PCR-amplified pROKA-putA with primer pair CRE-F/CRE-R (Table 2). The PCR product was treated by *Dpn*I, purified, phosphorylated with PNK (NEB), and subsequently self-ligated with T4 ligase at 16°C overnight. The resulting plasmid, pROKA-putA__CRE_, was inserted into *E. coli* DH5α (Table 1).

To replace the *putA* promoter sequence in pROKA-putA with that of *rocD*, we PCR-amplified pROKA-putA with primer pair promoter-Vec-F/Vec-R (Table 2). In contrast, the *rocD* promoter was PCR-amplified with primer pair PrD-F/PrD-R (Table 2). PCR products were assembled by the Gibson method (29), resulting in pROKA-putA__ProcD_ (Table 1). The plasmid was inserted into *E. coli* DH5α by transformation.

After verification of the correct sequence by DNA sequencing, the plasmids were electroporated into *S. aureus* RN4220 and subsequently transduced into *S. aureus* strain Newman with ϕ85.

To construct the *putA* expression plasmid pYJ335-putA, the *putA* gene, including 300-bp upstream sequence, was PCR-amplified with the primer pair P3356/P3357 (Table 2). The multi-copy plasmid pYJ335 was also PCR-amplified with the primer pair P3358/P3359(17) (Table 2). To make the pYJ335-argDCJB plasmid, the entire *argDCJB* operon, including 100-bp upstream sequence, was PCR-amplified with the primer pair P3418/P3419 (Table 2). The vector pYJ335 was PCR-amplified with the primer pair P3416/P3417(17) (Table 2). pYJ335-proC was constructed similarly. The *proC* gene was PCR-amplified with the primer pair P5173/P5174, whereas the vector pYJ335 was PCR-amplified with primer pair P5175/P 5176 (Table 2). The PCR products were treated with *Dpn*I (NEB, R0176) at 37°C for 30 minutes, purified with PCR purification kit (Qiagen), and assembled by Gibson method (29). The assembled DNA was inserted into *E. coli* DH5α by transformation. The insert DNAs of all plasmids were verified by DNA sequencing. Finally, the plasmids were electroporated into *S. aureus* RN4220 and subsequently transduced into *S. aureus* strain Newman with ϕ85.

### Construction of LacZ reporter plasmids

The *putA* promoter (P*putA*) and the *rocD* promoter (P*rocD*) were PCR-amplified with the primer pairs P3348/P3349 and P3352/P3353, respectively (Table 2). The plasmid vector sequence was PCR-amplified from pOS1-P2640-lacZ with the primer pairs P3350/P3351 and P3354/P3355 (Table 2). The PCR products were assembled by the Gibson method (29). The assembled plasmids were inserted into *E. coli*, purified, and the insert DNA sequence was confirmed by DNA sequencing. To generate cis-element deletion mutants of PputA, we PCR-amplified pOS1-PputA-lacZ plasmid with the reverse primer 3551 and one of six forward primers Δ1 ~ Δ6 (Table 2). The PCR products were treated with *Dpn*I at 37°C for 30 minutes, purified with a PCR purification kit (Qiagen). The purified DNA was phosphorylated with PNK (NEB), self-ligated with T4 ligase (NEB), and inserted into *E. coli* DH5a by transformation. The fidelity of insert DNA was verified by DNA sequencing for all resulting plasmids. The sequence-verified plasmids were electroporated into *S. aureus* RN4220 and subsequently transduced into *S. aureus* strain Newman with ϕ85.

### RNA extraction and quantitative reverse transcription-PCR (qRT-PCR)

Cells were grown in CDMG and harvested by centrifugation from bacterial cultures at 3 h post-inoculation for the mid-exponential growth phase and at 18 h post-inoculation for stationary growth phase cells. Total RNA was isolated from the harvested cultures using RNeasy mini prep kit (QIAGEN). Genomic DNA was eliminated with TURBO DNA-free™ kit (Invitrogen). The cDNA was synthesized from 2 μg of total RNA with SuperiorScript Master mix (Enzynomics). cDNA reaction mixture (8 μl) was used for PCR-amplification with SYBR green PCR reagent (Enzynomics) and the cognate primer pairs (Table 2). Reactions were performed in a MicroAmp Optical 96-well reaction plate (Applied Biosystems) and monitored with the 7300 Sequence Detector (Applied Biosystems). DNA was amplified 40 cycles at the following condition: 95°C for 10 sec, 60°C for 15 sec, 72°C for 30 sec. All qRT-PCR experiments were performed in triplicate and repeated independently three times. 16S rRNA was used as an internal control, and the results were analyzed by the comparative C_T_ method(30).

### β-galactosidase assay

The test strains were grown in CDM containing 15 μg/ml of chloramphenicol at 37°C for 16 h, and the optical density at 600 nm (OD_600_) was measured for the cultures. The bacterial cells were collected by centrifugation and suspended in Z-buffer (60 mM Na_2_HPO_4_, 40 mM NaH_2_PO_4_, 10 mM KCl, 1 mM MgSO_4_, 50 mM 2-mercaptoethanol, pH 7.0). The suspension was treated with lysostaphin (50 μg/ml) at 37°C for 30 min. After the addition of 4 volumes of Z-buffer to the lysed cells, 500 μl of the cell lysate was mixed with 10 μl of O-nitrophenyl-beta-galactopyranoside (ONPG, 12 mg/ml) and incubated at room temperature for 15 min - 1 h. When yellow color developed, 500 μl of Na_2_CO_3_ (1 M) was added, and the samples were centrifuged; then, the optical density of the supernatant was measured at 420 nm and 550 nm. The LacZ activity was determined by Miller units (U=1000×[(OD_420_)-(1.75×OD_550_)]/[(incubation time)×(volume)×(OD_600_)]). This assay was performed in triplicate and independently repeated at least three times.

### Isolation of *proC* transposon mutant

The mix of Phoenix transposon mutant library was grown in TSB containing 10 μg/ml erythromycin at 37°C overnight. Cells were collected from 1 ml culture by centrifugation, washed twice with 1 ml sterile water, and suspended in 1 ml sterile water. The cell suspension (100 μl) was spread on a CDMR agar medium without arginine (CDMR-R) and incubated at 37°C for 2 days. The resulting colonies were examined for their growth in CDMR-R broth. The transposon insertion site was determined by inverse PCR with the primers P142 and P143 (Table 2), as described previously(18).

### Quantification of pyrroline-5-carboxylate (P5C)

P5C was quantified by the ninhydrin derivatization method(19, 20). In the method, the test strains were grown in CDM broth at 37°C for 18 h. The bacterial cells were collected by centrifugation and suspended in 500 μl of TSM (50 mM Tris, 0.5 M sucrose, 10 mM MgCl_2_, pH 7.5). The suspension was treated with lysostaphin (50 μg/ml) at 37°C for 15 min and centrifuged. The protoplasts pellet was suspended in 1 ml of 50 mM Tris (pH 8.0) and mixed with 0.25 ml of perchloric acid (3.6 N), 0.25 ml of ninhydrin (2% w/v). The mixture was boiled for 15 min, and the insoluble ninhydrin derivatives were collected by centrifugation. The collected derivatives were dissolved in 0.5 ml of ethanol, and 0.5 ml of 100 mM Tris (pH 8.0) was added. After centrifugation, the supernatant was collected, and its optical density was measured at 620 nm. The sample’s optical density was normalized by the culture optical density at 600 nm. Finally, the P5C concentration was calculated with the molar extinction coefficient (1.96×10^5^) and normalized by OD_600_ of the culture (P5C concentration=OD_620_/[ε/(volume)×(OD_600_)]). This assay was performed in triplicate and independently repeated at least three times.

### Generation of PutA antibody

To produce the PutA protein, we generated pET28a-putA. pET28a was amplified with primers P4105/P4106. And *putA* gene was amplified with primers 4910/4911. The purified PCR products were assembled by Gibson method(29) and inserted into *E. coli* DH5α. After confirmation of the *putA* sequence by DNA sequencing, the plasmid was inserted into *E. coli* BL21(DE3).

To express His-tagged PutA (His-PutA), we grew *E. coli* BL21(DE3) carrying pET28a-putA in LB at 37°C and treated with 1 mM IPTG (Isopropyl β-D-1-thiogalactopyranoside). After 3 h incubation, the cells were collected, and His-PutA was purified with Ni-column chromatography (Qiagen). The purified His-PutA protein (50 μg) was 1:1 (v/v) mixed with either complete (1^st^ injection) or incomplete (2^nd^ and 3^rd^ injection) Freund’s adjuvant (Sigma) and injected into C57BL/6 mice (6 – 8 week old) intramuscularly in two-week intervals. One week after the 3^rd^ injection, all mice were anesthetized, and blood was collected via cardiac puncture.

### Western blot analysis of PutA

The test strains were grown in CDM broth at 37°C and harvested at exponential growth phase, and normalized to OD_600_ = 4. The cells were collected by centrifugation and suspended in 100 μl of TSM, and treated with lysostaphin (50 μg/ml) at 37°C for 15 min. The protoplasts cells were collected by centrifugation and suspended in 80 μl of lysis buffer (50 mM Tris, 150 mM NaCl pH 7.4). The lysed cells were added 1 ul of DNaseI and incubated at 37°C for 30 min. Twenty microliters of 5× SDS sample buffer were added to the cell lysates, and samples were boiled for 10 min. After brief centrifugation, 10 μl of samples were subjected to 12% SDS-PAGE. Western blotting was carried out with the PutA antibody (1:1000 dilution). Signals were detected by SuperSignal West atto chemiluminescent substrate (Thermo Scientific).

## Acknowledgments

We thank Suzan Rooijakkers at the University Medical Center Utrecht, the Netherlands for sharing pKOR1-mcs. This work was supported by the NIH fund (AI143792) to TB and by the Korea Health Technology R&D Project through the Korea Health Industry Development Institute (KHIDI), funded by the Ministry of Health and Welfare, Republic of Korea (grant number HI19C1095) to BJ. The funders had no role in study design, data collection, and interpretation or the decision to submit this work for publication.

